# Auto-encoded Latent Representations of White Matter Streamlines for Quantitative Distance Analysis

**DOI:** 10.1101/2021.10.06.463445

**Authors:** Shenjun Zhong, Zhaolin Chen, Gary Egan

## Abstract

Parcellation of whole brain tractograms is a critical step to study brain white matter structures and connectivity patterns. The existing methods based on supervised classification of streamlines into predefined streamline bundle types are not designed to explore sub-bundle structures, and methods with manually designed features are expensive to compute streamline-wise similarities. To resolve these issues, we propose a novel atlas-free method that learns a latent space using a deep recurrent auto-encoder. The method efficiently embeds any length of streamlines to fixed-size feature vectors, named streamline embedding, for tractogram parcellation using unsupervised clustering in the latent space. The method was evaluated on the ISMRM 2015 tractography challenge dataset with discrimination of major bundles using unsupervised clustering and streamline querying based on similarity. The learnt latent streamline and bundle representations open the possibility of quantitative studies of arbitrary granularity of sub-bundle structures using generic data mining techniques.

## 1. Introduction

Diffusion tractography is a post-processing technique that generates streamlines as a proxy to study the underlying white matter fiber networks. Diffusion-weighted imaging (DWI) is a magnetic resonance imaging (MRI) technique that measures the random motion of water molecules in biological tissues (Jones, 2010). In brain imaging, DWI is used to characterize the distribution and orientation of the underlying white matter pathways. DWI data provides quantitative diffusion measurements, including the apparent diffusion coefficient (ADC) and fractional anisotropy (FA), that can be further analysed to model the voxel-wise magnitude and orientation information. Current techniques include diffusion tensor imaging (DTI) and spherical harmonics based methods (e.g. constrained spherical deconvolution (Tournier et al., 2008)) for high angular resolution diffusion imaging data (HARDI). Together with fiber tractography, DWI data enable investigation of brain network connectivity using three dimensional (3D) reconstruction of streamlines to represent white matter fibre pathways in the brain.

Streamlines consist of a series of 3D spatial positions that can be grouped into meaningful bundles using streamlining clustering or segmentation algorithms. Since manual streamline anatomical labelling is extremely time-consuming, many automated or semi-automated labelling solutions have been proposed to identify and label streamlines to pre-defined anatomical white matter fiber bundles, e.g. the corticospinal tract (CST) and corpus callosum (CC). The automated solutions to the streamline labeling problem can be: (i) region-of-interest (ROI) based filtering, i.e. the assignment of a streamline is determined by defining ROIs where the streamlines should pass, bypass and terminate (Jones & Pierpaoli, 2005; Wakana et al., 2007; Cammoun et al., 2012; Colon-Perez et al., 2016; Yendiki et al., 2011; Wassermann et al., 2016); (ii) distance based methods that label a streamline to its spatially closest reference streamline tract (Maddah et al., 2005; Clayden et al., 2007; Labra et al., 2017; Corouge et al., 2004); and (iii) classifier-based methods that learn a model from annotated streamline data to correspond to a streamline pre-defined bundle class, with deep learning (DL) based classifiers with convolutional neural networks (CNNs) (Gupta et al., 2017; Gupta et al., 2018; Ugurlu et al., 2018; Zhang et al., 2020) and graph convolutional networks(GCNs) (Liu et al., 2019) used for supervised learning. The ROI-based and distance-based approaches require carefully defined ROIs and reference tracts, and are difficult to adapt to group studies, due to inter-subject variability and time consuming manual annotation. The classifier based approaches require a large number of labeled streamlines to train the DL models, and are only limited to predict a fixed number of predefined streamline bundles according to the manual annotations for model training in the dataset.

Supervised learning approaches are limited to predefined streamline bundle types and require a large number of annotated streamlines via manual or semi-automated annotations. Conversely, unsupervised learning methods are able to cluster streamlines in various granularities without knowledge of the labels. Early methods focused on the design of spatial distance functions. The mean closest point distance (Corouge et al., 2004) was widely used to compute a distance matrix of all the streamline pairs, i.e. inter-streamline distances, and conduct clustering algorithms to decompose streamlines into plausible bundles, including spectral clustering (Ziyan et al., 2009) and hierarchical clustering (Zhang et al., 2008; Jianu et al., 2009). The method calculates the Euclidean distances of each point on a streamline to all points in another streamline. A more computational efficient approach, QuickBundles (Garyfallidis et al., 2012), uses the minimum average direct-flip (MDF) distance, and the element-wise distance measurements of two streamlines that are required to have the same length. The method does not handle different streamline lengths and spatially various endpoints where streamlines belong to the same bundle types but have endpoints that are not apart from each other.

The efficiency of streamline clustering can be improved using a feature space which encodes the streamlines into corresponding feature representations, termed streamline embedding. Measuring the distances between a streamline and the reference streamlines (i.e. the landmarks) can capture the relative position and shape information of a streamline. By selecting a predefined number of reference streamlines, a distance-based feature vector can be constructed with the same length. Olivetti et al. (2012) designed three different policies to select reference streamlines, and used a symmetric minimum average distance (Zhang et al., 2008) to encode each streamline to a feature vector. Berto et al. (2021) selected reference landmarks from global and local perspectives, and considered the distances of the streamline endpoints to the global reference landmarks and the predefined ROIs. Lam and co-workers (Lam et al., 2018) also included curvature and torsion as features to represent a streamline. However, methods that rely on landmark selections assume all streamlines including the reference landmarks, have the same length and the embedding performance is related to the number of reference landmarks.

The shape and position of streamlines can be directly modelled by manually designing the feature scheme. Brun et al. (2004) captured the position, shape and connectivity information of streamlines by constructing a 9D vector consisting of the 3D mean vector and the lower triangular part of the covariance matrix of the streamline points. In the work from Maddah et al. (2006), the coefficients of the 3D quintic B-spline were used as the feature vector representation of a streamline. Batchelor et al. (2006) and Chung et al. (2010) used Fourier series as the basis function and the shape of streamlines were represented using a series of Fourier coefficients. This representation model was adopted by Wu et al. (2020) with dictionary prototypes learnt for each predefined bundle type and streamlines classified based on the minimized sparse coding residuals. In the work of FiberMap (Zhang et al., 2020), the sequential information of a streamline was converted to a 2D feature map by splitting the coordinates in the x, y and z axis as three channels, flipping and repeating the spatial coordinates. However, the FiberMap representations of streamlines lack a distance measurement definition so cannot be directly used for quantitative analysis, and the streamlines are assumed to have the same length. Nevertheless, the 2D feature map can be with 2D CNN for supervised learning tasks.

Embedding or representation in a latent space is an effective and robust method for data-driven learning. A number of basis functions have been used to represent streamlines as combinations of the basis functions. Continuous basis functions can be learned with dictionary learning techniques and streamlines can be represented as a combination of dictionaries, to encode streamlines as fixed-size coefficients that represent the feature vector (Alexandroni et al., 2017; Kumar et al., 2019). Similar to our previous work (Zhong et al., 2020), Legarreta et al. (2021) used a 1D CNN-based auto-encoder to learn a latent space to measure distances from the latent streamline representations to manually selected reference streamlines. However, convolutional neural networks do not capture the sequential nature of streamlines and require fixed size inputs, which limits the ability of the neural network to handle variable streamline lengths.

In this paper, we extended our previous work (Zhong et al., 2020), and proposed and validated a novel recurrent neural network (RNN) to learn latent streamline representations. The key idea is motivated by the successful applications of sequential modelling in natural language processing (NLP) where recurrent neural networks perform well on capturing long sequential information and are proven to effectively model latent features (Mikolov, Sutskever, et al., 2013; Cho et al., 2014; Mikolov, Chen, et al., 2013; Le & Mikolov, 2014). This work uses a Long Short Term Memory (LSTM) RNN to auto-encode any length of streamlines into fixed-size latent vectors. Similarities are measured in the latent space by simply computing the Euclidean distances between the streamlines. Compared to the existing methods, our method can handle streamlines of any length without resampling to ensure equal streamline lengths, and capture the sequential information of streamlines by the latent representation. Furthermore, we propose an effective way to construct group or streamline bundle embedding via averaging latent vectors. Embedding the streamlines and bundles enables clustering, querying, filtering and downstream quantitative analysis of the white matter streamlines.

## 2. Methods

In this section, we present the proposed neural network architecture and training scheme, introduce the NN-based streamline embedding mechanism to convert any length of streamline to a multi-dimensional feature vector, show the effective way to obtain embedded streamline bundling from the streamline embedding, and describe the experiments and corresponding validation processes.

### 2.1 Recurrent Autoencoder

Long Short Term Memory (LSTM) is a type of recurrent neural network (RNN) unit that encodes sequential information from variable length data samples (Hochreiter & Schmidhuber, 1997). It consists of several gates, and output a hidden state *h* and an output *o*. At each time step*t*, the hidden state *h*(*t*)is updated by feeding the corresponding inputs and the state from the previous time step *h*(*t*−1), as

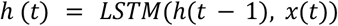

where *x*(*t*)is the input data of the current time step. Models with LSTM can learn to predict the next element in a sequence, by fitting the probability distribution, *p*(*x*_*t*_ | *x*_*t*−1_, *x*_*t*−2′_, …,*x*_1_) over a sequence. Therefore, LSTM has been widely used in areas where the data have a sequential temporal component, e.g. encoding video features (Bin et al., 2018; Gao et al., 2017), natural language processing (Cho et al., 2014; Sundermeyer et al., 2012) and brain functional analysis(Oota et al., 2019).

Due to the temporal memorization property, LSTM units are successfully used in encoder-decoder neural networks to map an input sequence to a target sequence, which is commonly known as the Seq2Seq auto-encoder network (Sutskever et al., 2014). The encoder transforms the input sequence into a fixed-length latent vector, by feeding each element of an input sequence one by one and updating the hidden states where the last hidden state is considered to contain the summarized information of the entire input sequence. On the other hand, the decoder takes the summary vector from the encoder and generates elements sequentially. During training, encoder-decoder networks are trained in an end-to-end manner, while the trained encoder and decoder can be used independently or together during inference. The encoder can be used to embed an input sequence into a latent vector alone.

### 2.2 Network Architecture and Training

As shown in Fig 1, a LSTM-based seq2seq auto-encoder was used to embed the streamlines into a fixed length latent representation. The encoder received the first half of each streamline and output a feature vector; and the decoder was trained to predict the second half of the same streamline during the training. The encoder and decoder networks both contained a single layer of LSTM, with 128 hidden dimensions. We used Adam optimizer to minimize the loss (i.e. Mean Square Loss) between the predicted and the ground truth, using a learning rate of 1e-3 and a batch size of 128.

**Fig 1.**
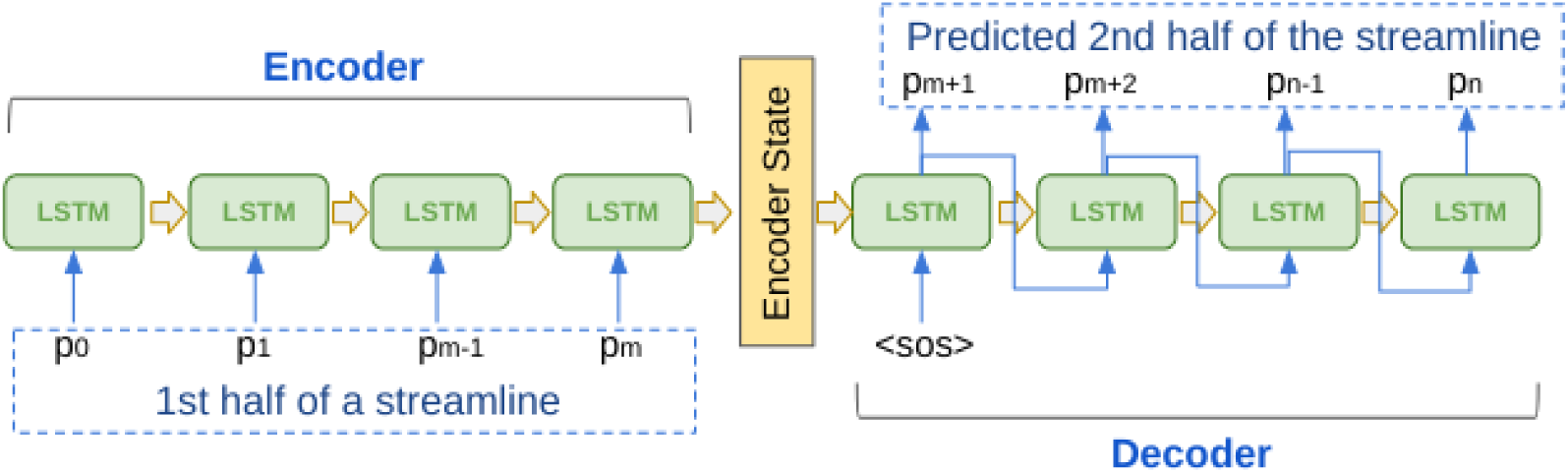
LSTM-based auto-encoder architecture. The first half of the streamline was fed into the encoder of the network sequentially and the hidden state of the encoder was passed to the decoder. The decoder learned to point-by-point predict the second half of the streamline.

**Fig 2.**
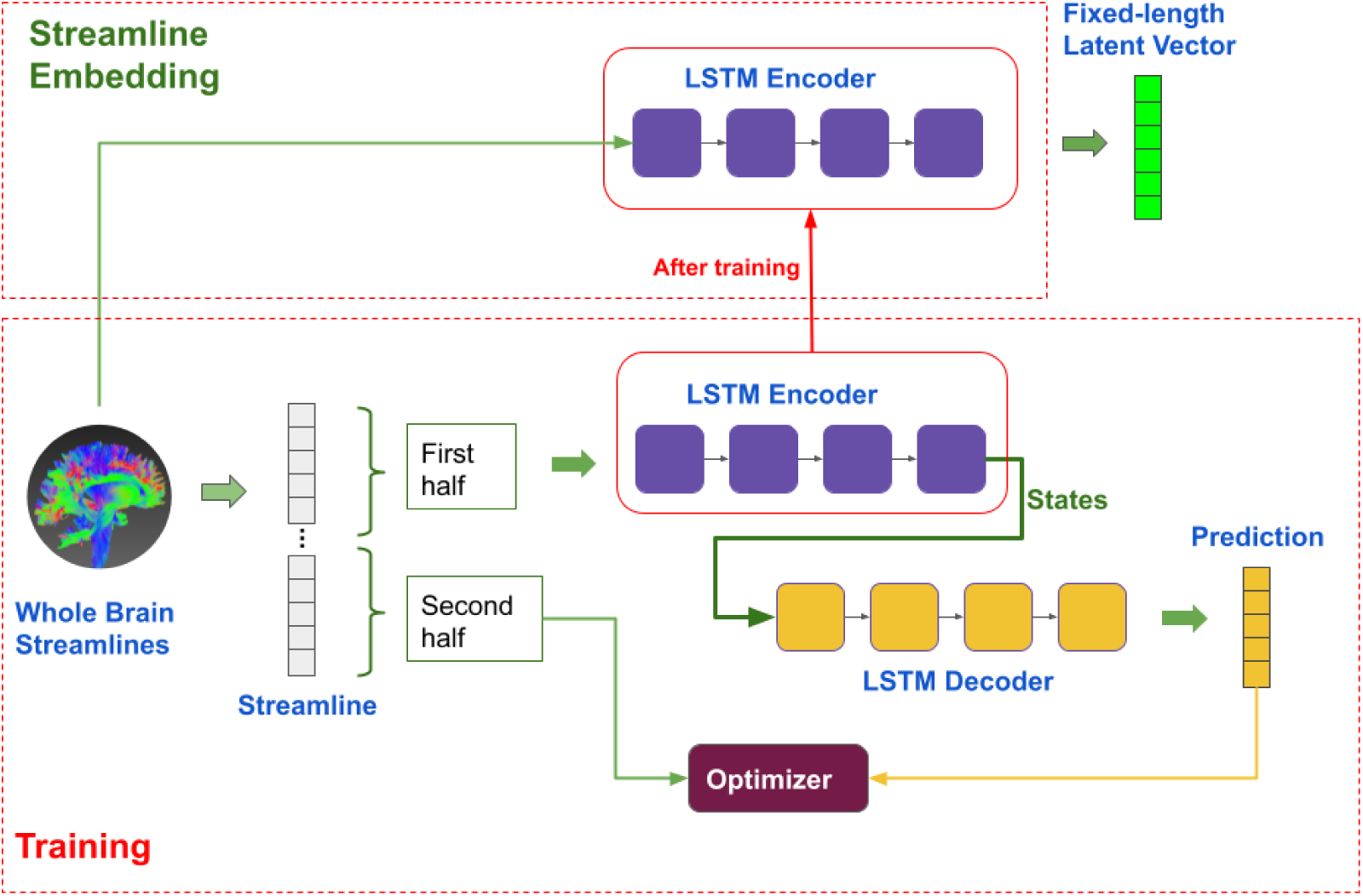
Training and validation workflow. During training, the whole brain streamlines were used to train the auto-encoder. After trained, only the encoder part was used for embedding any length of streamlines into their fixed-size latent vectors.

For each training iteration, the input sequence (i.e. the first half of a streamline) was appended with a special token, (0,0,0) to indicate the end of the sequence and the same token was appended to the start of the output sequence (i.e. the second half of the same streamline). The training samples in a batch were padded with zeros to match the length of the longest streamline in the batch. The LSTM hidden and cell states computed from the encoder were passed to the decoder. During decoding, a special token was fed, with value, (0,0,0) to indicate the start of the decoding process, then the decoding loop predicted the next position iteratively. No teacher forcing technique was used, meaning that the input to predict the next position during decoding was the LSTM output from the previous decoding step. The loss was computed by aggregating the individual losses between each predicted position of the second half streamline and the ground truth. A gradient clip of 1.0 was used to prevent gradient explosion, and the training samples were randomly shuffled between each epoch.

### 2.3 Streamline Embedding

After the model was trained, the encoder was used to embed the streamlines into fixed-length latent vectors, i.e. the streamline length was converted into the corresponding latent representation. The length of the latent representation could be configured during model training, by setting the hidden dimension in the LSTM cell, in this case, we used a 128 hidden dimension which resulted in the latent vector to be the length of 128. A larger latent dimension, e.g. 256 or 512, leads to a larger parameter space for the auto-encoder, and consequently more computational resources during training and inference. On the other hand, a small latent space, e.g. 32 or 64, requires less computing power, but it is prone to errors if the model is under-fitted.

### 2.4 Streamline Bundle Embedding

The embedding of each streamline bundle can be formed to represent streamline groups, by averaging the latent representations of the streamlines in the bundle or group. The bundle embedding, *B* was defined as the explicit average of the streamline embeddings, *S*, that belonged to that bundle, i.e.

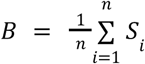

where *n* is the number of streamlines in a bundle. In this way, the embedded bundle had the same vector length as the streamlines, which enabled similarity measurements to be computed between the streamline and bundles.

### 2.5 Similarity measurement and streamline clustering

The similarity between two streamlines can be directly measured by calculating the Euclidean distance of their corresponding high-dimensional latent vectors. Similar to the application in NLP where semantically similar words, sentences or paragraphs are closer to each other, the auto-encoded streamlines have the property that anatomically similar streamlines are located closer in the latent space. Hence, the streamline-wise similarity can be calculated using the Euclidean distance metric, which has the computing complexity of O(n). The complexity is only dependent on the length of the latent vector which is customizable when training the auto-encoder, and a smaller hidden dimension can be used, for example 32 or 64, to reduce the distance metric computation.

Similarly, bundle-wise distance and similarity can be measured and any streamline can be classified to defined bundles by the sorted distances. As explained in section 2.4, streamline bundle embedding can be represented as the average of all or a subset of the embedded streamlines. The similarity between different bundles can be calculated with the Euclidean distance and the distance between a streamline and a bundle can be computed using their corresponding embedded vectors. In this way, if the bundles are determined, any streamlines can be easily classified to the bundle types by simply measuring the distances.

The latent embedding enables streamline clustering with any clustering algorithm in an unsupervised manner. Each streamline is embedded to a feature vector using the trained encoder, and can be treated as delimited data where each row represents a streamline and columns are the latent features. Since the embedding part is a separate step, any clustering algorithms can be used to segment the streamlines into groups. In our work, the k-mean algorithm was used to cluster streamlines into anatomically plausible groups.

### 2.6 Dataset

The dataset used in the study was from the ISMRM 2015 Tractography Challenge (Maier-Hein et al., 2015) which contained 25 predefined types of streamlines in the TCK format. There were in total 200,432 streamlines available in the dataset, which covered 25 different types, such as corpus callosum (CC), left and right superior cerebellar peduncle (SCP). The streamlines were further split into training (80%) and validation (20%) datasets. The model with the minimal loss value was selected for the subsequent experiments, where all of the streamlines were used.

### 2.7 Experiments and Validation

To validate the latent representations that capture useful information about the streamlines, several streamline clustering experiments were conducted. Streamlines were clustered in the latent space, using the basic K-means algorithm, and assigned to k clusters where k was customizable. In our work, the experiments included: (i) segment symmetric bundles, e.g. left and right SCP, left and right ICP, anterior and posterior commissures, and tests whether the model can separate left/right or anterior/posterior counterparts; (ii) dimension reduction of latent vectors into 2D using t-SNE algorithm and visual check that the features were separable even in lower dimension; (iii) decomposition of the bundles on the left hemisphere (excluding CC, fornix and middle cerebellar peduncle (MCP)) into ten clusters and validation that the clustered streamlines could be mapped back to the ground truth bundle types; (iv) query and filtering of streamlines based on distance measurements; and (v) validation of the streamline classification accuracy by sorting of the distances in the latent space. The clustered streamlines were further converted back and stored as TCK files for visualization and subsequent processing tasks.

The classification accuracy was assessed by computing the top-k metrics, which were defined as the percentage of correctly assigned streamlines based on the Euclidean distances to the embedded bundles. The top-1, top-3 and top-5 were recorded to examine the performance of the learnt latent representation model.

## 3. Results

The model converged after training for typically two epochs. The best performed model with the lowest validation loss on the validation data was chosen for reporting the results. In the inference phase, only the encoder was used for embedding streamlines.

### 3.1 Streamline Clustering

The latent features can distinguish spatially symmetric structures with unsupervised clustering algorithms. As shown in Fig 3(a-c, e-g) and Fig 4, the streamline bundles that have symmetric parts (left and right), e.g. uncinate fasciculus (UF), inferior longitudinal fasciculus (ILF), parieto-occipital pontine tract (POPT), superior longitudinal fasciculus (SLF), corticospinal tract (CST) and optic radiation (OR) were perfectly clustered, while the results for the fronto-pontine tracts (FPT) were correct with minor bias. However, the left and right components of the inferior cerebellar peduncle (ICP), superior cerebellar peduncle (SCP) and cingulum were not separated. Fig 3(d) and 3(h) show the feature distributions of the anterior commissure (CA) and posterior commissure (CP) and the left/right CST after dimension reduction from 128 to 2. These bundles were clearly separable indicating that the learned latent representations captured the necessary spatial information of the streamlines.

**Fig 3.**
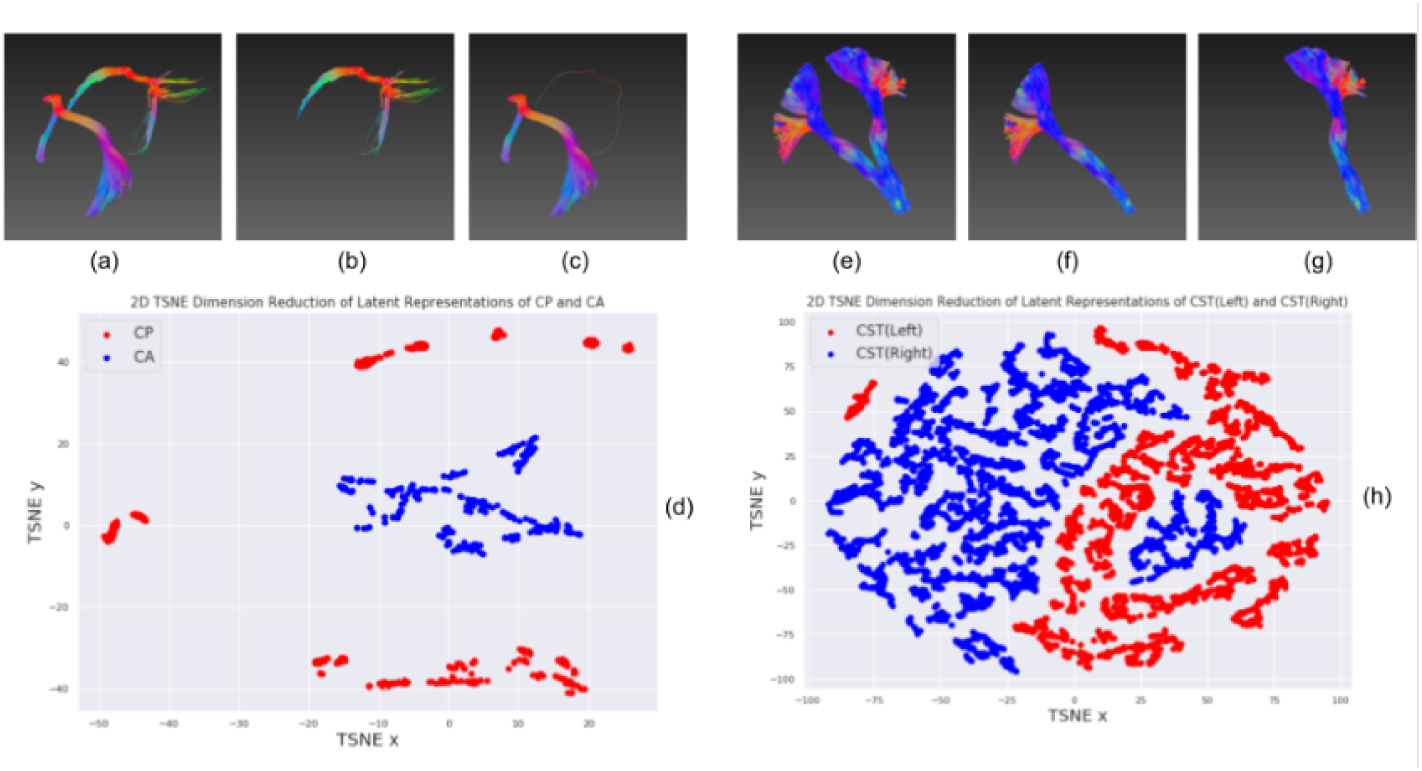
Streamline clustering of CA/CP and left/right CST, and their t-SNE 2D feature distributions. (a-c) Visual inspection of the clustering results for CA and CP; (d) the t-SNE 2D projection of the latent vectors of all streamlines in CA and CP showing a clearly separation; (e-g) clustering results of the left and right CST; and (h) the t-SNE 2D projection of the latent vectors of all streamlines for the left and right CST.

**Fig 4.**
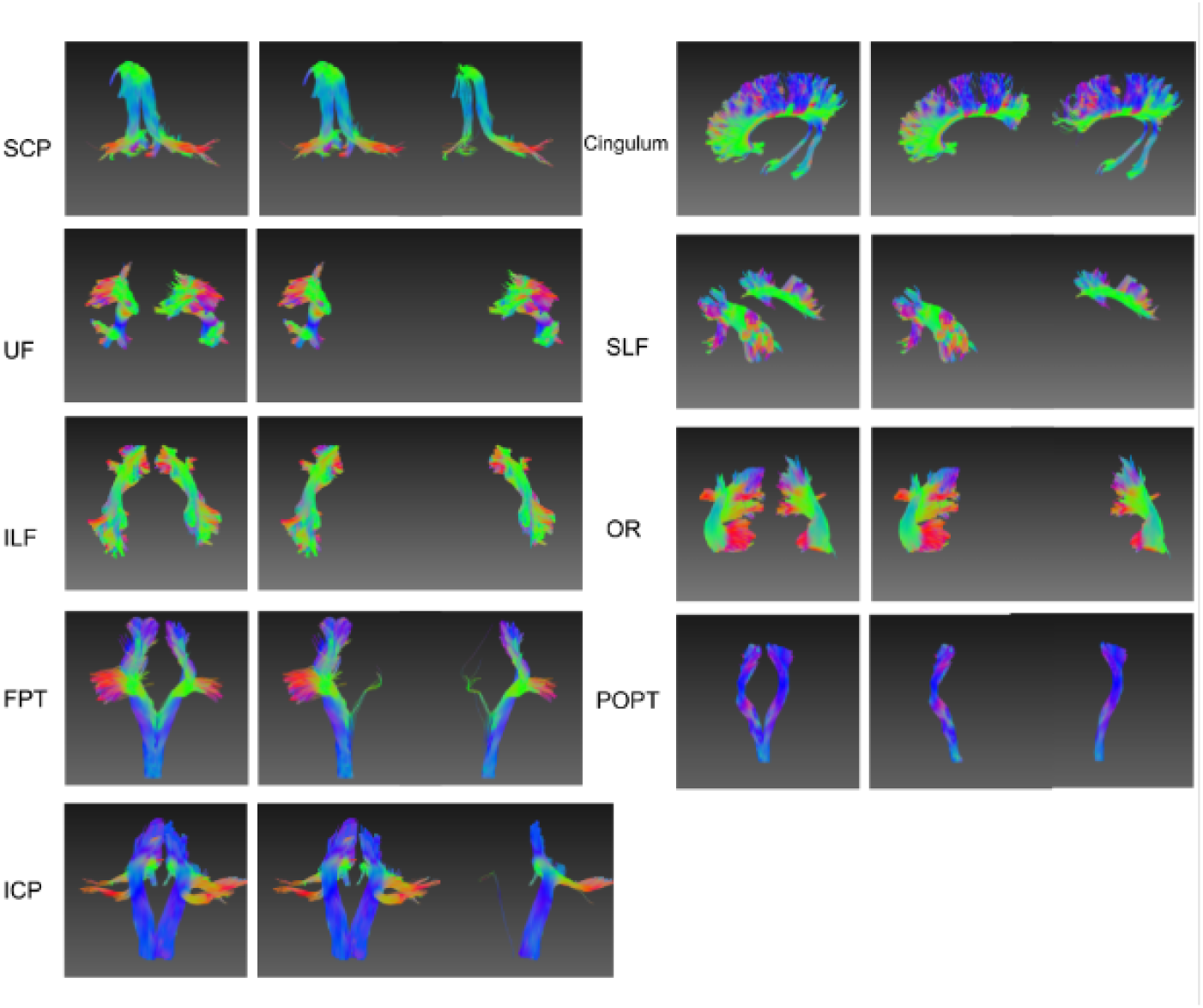
Streamline clustering for left/right symmetric bundles. The symmetric bundles are embedded into latent representations and clustered into two groups. UF, ILF, POPT, SLF, CST and OR are perfectly clustered, while the ICP, cingulum and SCP failed to discriminate between symmetric parts.

A more challenging task was decomposition of the streamlines in the entire left hemisphere into meaningful bundles. In this case, the corpus callosum (CC), fornix and middle cerebellar peduncle (MCP) were excluded. As shown in Fig 5, after clustering into ten classes, the model produced major bundles with minor bias, including for the UF, CST, OR, SLF and cingulum. Some of the bundles overlapped, including the SCP and ICP, POPT and FPT, and ILF and OR. Fig 6 shows the 2D feature embedding of the ten bundles from the left hemisphere after dimension reduction with t-distributed stochastic neighbor embedding (t-SNE) (Van der Maaten & Hinton, 2008). The original feature vectors of 128 dimensions were reduced to two dimensions for visualization purposes. Interestingly, streamlines of the same bundle type were visualized closer to each other and separable from other types, even in a two dimensional space representation.

**Fig 5.**
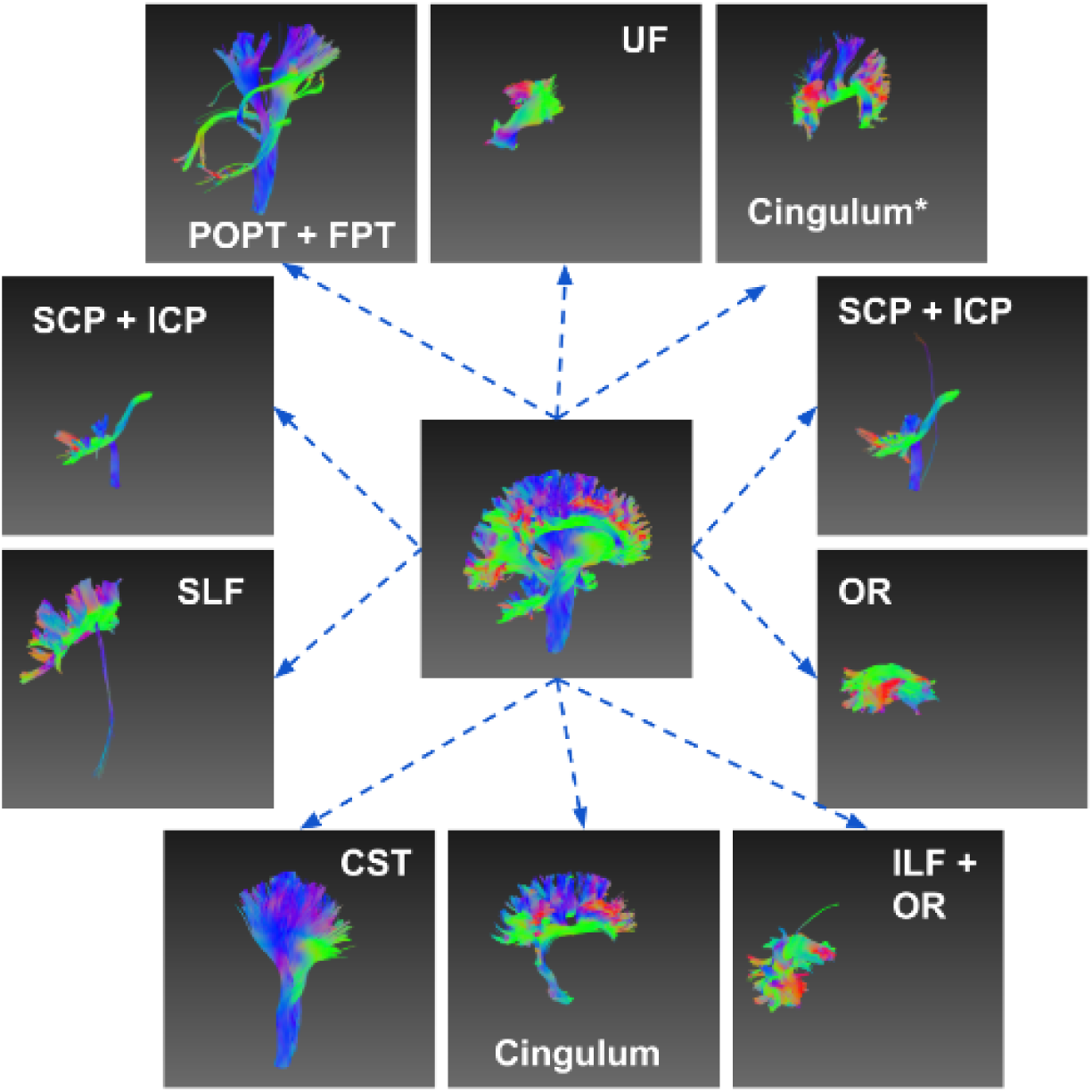
Decomposition of the left hemisphere and the clustered bundles. All the streamlines in the left hemisphere were converted into latent vectors and clustered into ten groups. Most of the streamline bundles can be extracted, while some groups contain parts from two bundles.

**Fig 6.**
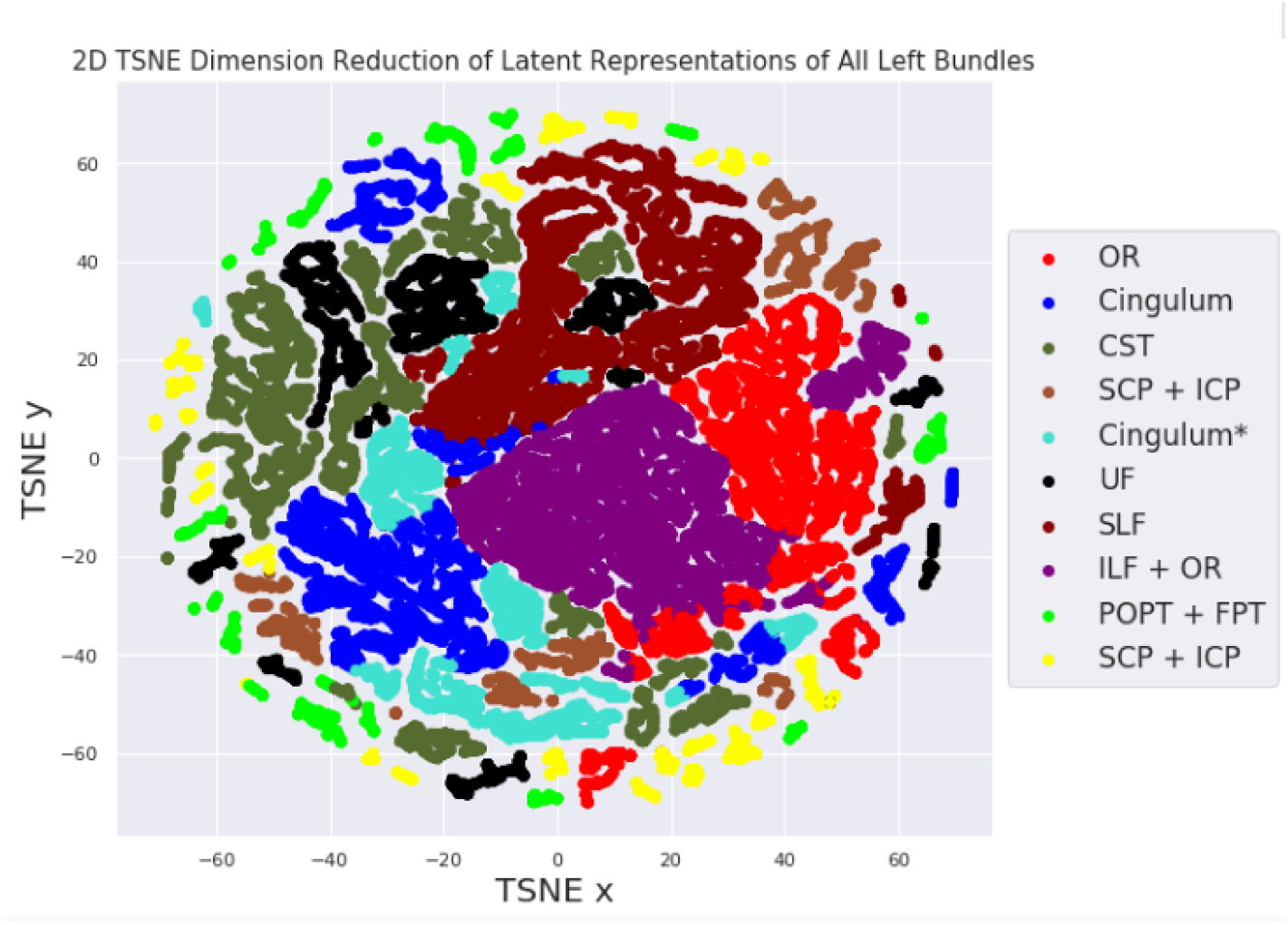
t-SNE 2D feature distributions of the ten selected bundles on the left hemisphere. The original feature vectors are 128 dimensions, and the dimensions are reduced to 2D by using t-SNE. Visual inspection of the feature distributions indicates that the streamlines from the same bundle types are closer to each other.

### 3.2 Streamline Classification

Using the synthetic dataset we measured the classification accuracy based on the sorted Euclidean distances between the streamlines and the 25 bundles. In Table 2 the results are reported for the top-1, top-3 and top-5, i.e. the correct label in the k nearest, where *k* ={1,3,5}. The overall top-1, top-3 and top-5 accuracy were 81.1%, 97.0% and 98.7%, i.e. around 81% of the time, the nearest bundle type is the correct type. The details of the classification accuracy for each bundle type can be seen in Table 1. Seven out of 25 bundles had classification accuracy of > 98% and mostly > 99%, including for the left and right SLF, UF, and ICP and for the left CST. However, the accuracy was poorer for the bundles in the top-1 metric, e.g. CC (54%), fornix (61%), left POPT (55%), left SCP (43%), right FPT (55%) and left cingulum (69%). For the top-3 metric all of the bundle types, excluding the CC and left FPT, achieved > 90% accuracy and twelve bundle types (out of 25) had a top-3 metric beyond 99%.

**Table 1.** Top K accuracy (out of 25) based on distance measurements for each bundle type.

|  |  |  |  |  |  |  |  |
| --- | --- | --- | --- | --- | --- | --- | --- |
| **Bundle** | **Top1** | **Top3** | **Top5** | **Bundle** | **Top1** | **Top3** | **Top5** |
| **OR_left** | 0.8875 | 0.9998 | 0.9998 | **CST_left** | 0.9979 | 0.9982 | 0.9982 |
| **MCP** | 0.9447 | 0.9999 | 0.9999 | **SLF_left** | 0.9989 | 0.9995 | 0.9995 |
| **FPT_left** | 0.6982 | 0.9652 | 0.9710 | **SCP_right** | 0.7699 | 0.9878 | 0.9994 |
| **Fornix** | 0.6197 | 0.8932 | 0.9862 | **FPT_right** | 0.5571 | 0.8254 | 0.9146 |
| **UF_right** | 0.9976 | 0.9981 | 0.9997 | **Cingulum_right** | 0.8730 | 0.9836 | 0.9949 |
| **CC** | 0.5440 | 0.8598 | 0.9290 | **Cingulum_left** | 0.6906 | 0.9450 | 0.9710 |
| **ILF_right** | 0.7613 | 0.9851 | 0.9934 | **CST_right** | 0.9629 | 0.9773 | 0.9979 |
| **POPT_left** | 0.5522 | 0.9706 | 0.9848 | **UF_left** | 0.9919 | 0.9919 | 1.0000 |
| **ILF_left** | 0.7310 | 0.9982 | 1.0000 | **CA** | 0.9002 | 0.9954 | 1.0000 |
| **POPT_right** | 0.8140 | 0.9152 | 0.9380 | **SLF_right** | 0.9977 | 0.9999 | 1.0000 |
| **ICP_right** | 0.9997 | 0.9997 | 0.9997 | **CP** | 0.7452 | 0.9781 | 0.9973 |
| **SCP_left** | 0.4368 | 0.9733 | 1.0000 | **ICP_left** | 0.9879 | 1.0000 | 1.0000 |
| **OR_right** | 0.8090 | 0.9999 | 0.9999 |  |  |  |  |

**Table 2.** Sorted embedding distances of a randomly sampled streamline.

| **Bundle** | **Distance** | **Bundle** | **Distance** | **Bundle** | **Distance** |
| --- | --- | --- | --- | --- | --- |
| **CST_right** | 0.719 | **Cingulum_right** | 2.755 | **FPT_left** | 3.927 |
| **UF_right** | 1.950 | **SCP_right** | 2.784 | **ICP_left** | 4.041 |
| **ILF_right** | 2.054 | **ICP_right** | 3.094 | **POPT_left** | 4.193 |
| **CP** | 2.082 | **MCP** | 3.172 | **ILF_left** | 4.219 |
| **Fornix** | 2.313 | **SCP_left** | 3.261 | **UF_left** | 4.771 |
| **FPT_right** | 2.390 | **CST_left** | 3.332 |  |  |
| **OR_right** | 2.414 | **Cingulum_left** | 3.486 |  |  |
| **POPT_right** | 2.578 | **SLF_left** | 3.723 |  |  |
| **SLF_right** | 2.599 | **OR_left** | 3.782 |  |  |
| **CC** | 2.727 | **CA** | 3.807 |  |  |

### 3.3 Streamline Querying

The results of querying similar streamlines based on different distance thresholds is shown in Fig 7. A streamline was randomly selected from the whole brain tractograms and treated as the seeding streamline. In Fig 7 the seeding streamline was from the right CST bundle. By measuring the distances between the seeding streamline and all other streamlines in the tractogram enables streamlines to be queried within different distance ranges. In this case, small distance thresholds, e.g. 0.125 and 0.25 led to small streamline bundles which were highly similar to the seeding streamline and were a subset of the right CST bundle. Increasing the distance thresholds resulted in the inclusion of more streamlines, with a threshold of 0.75 including the entire right CST bundle. Larger thresholds did not query additional streamlines since there were no streamlines within the distance range of 0.75 and 1. In practice, much smaller step sizes can be used to progressively explore the sub-structure of the white matter bundles.

**Fig 7.**
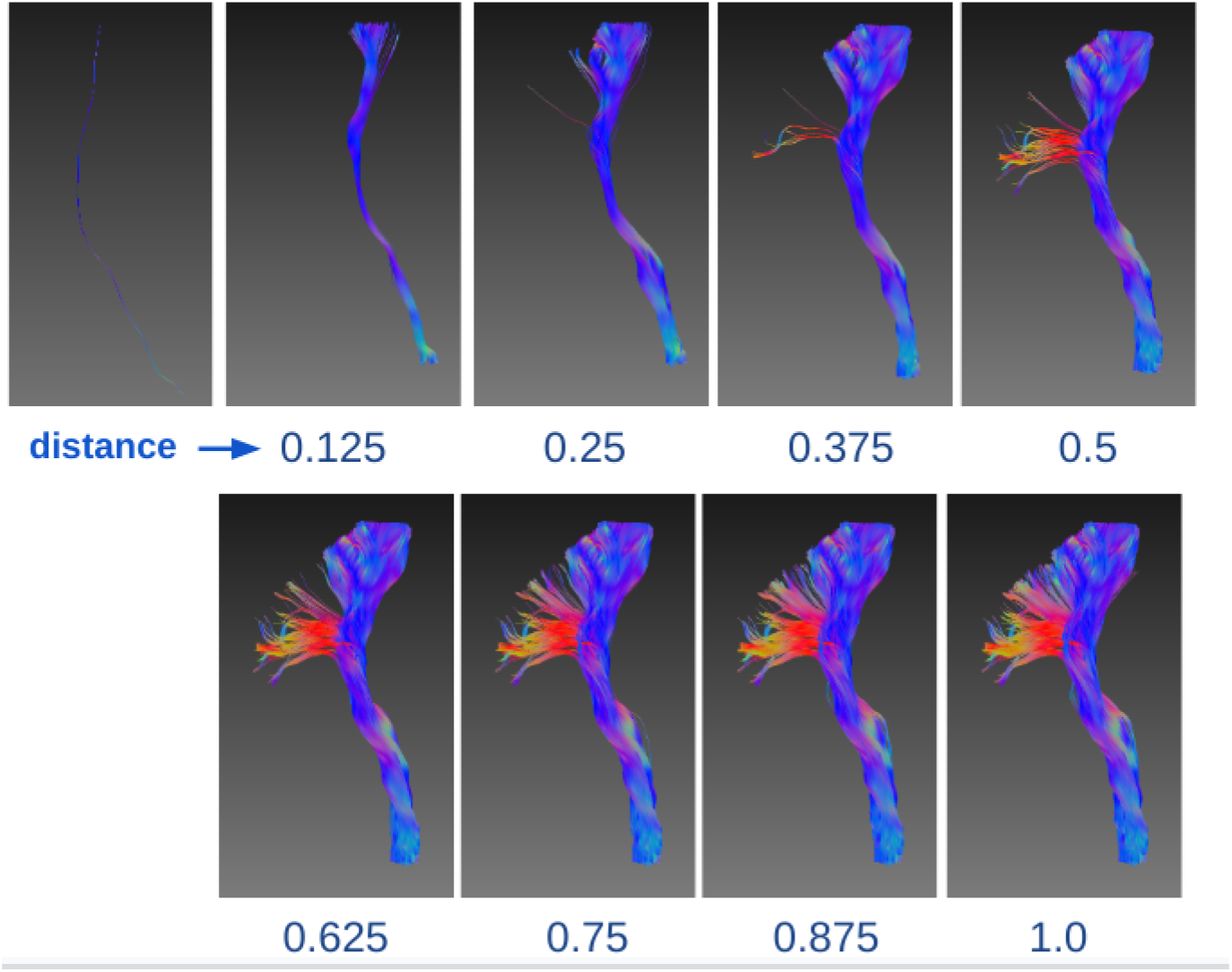
Query similar streamlines by various distances. A streamline is randomly selected from left CST and used as the seeding streamline to query similar streamlines based on various distance thresholds. Use of a smaller threshold results in sub-bundles that are very similar to the seeding streamline, while a larger threshold leads to the extraction of the entire left CST bundle.

Quantitatively, selecting the threshold can be guided by the distance measurements between the seeding streamline to the embedded bundles. Table 2 shows the streamline-bundle distances calculated for the same seeding streamline used in Fig 7. The distance between the seeding streamline and the bundle embedding of the right CST was 0.719, consistent with the observation that the threshold of 0.75 returns the streamlines from the entire right CST bundle. The distance to the second closest bundle type, the right UF bundle was 1.95. Hence, any threshold selection below that value did not include streamlines from other bundle types.

### 3.4 Bundle-wise Distance Analysis

Group-wise similarity between the embedded bundles can be analyzed by the distance matrix. Fig 8 shows the distances between the 25 predefined bundles in the synthetic dataset, using bundle embedding. Symmetric bundles, such as left and right ICP were closer to each other in the latent space, and the same property was observed for SCP and the cingulum indicating that the shape information was encoded by the neural network. SCP and ICP bundles were closer to each other, as part of the streamlines have similar spatial properties. Groups of bundles are spatially closer to each other in the top-left part of the distance matrix, including the CA, CC, CP, left and right CST, left and right cingulum, left and right FPT and the fornix.

**Fig 8.**
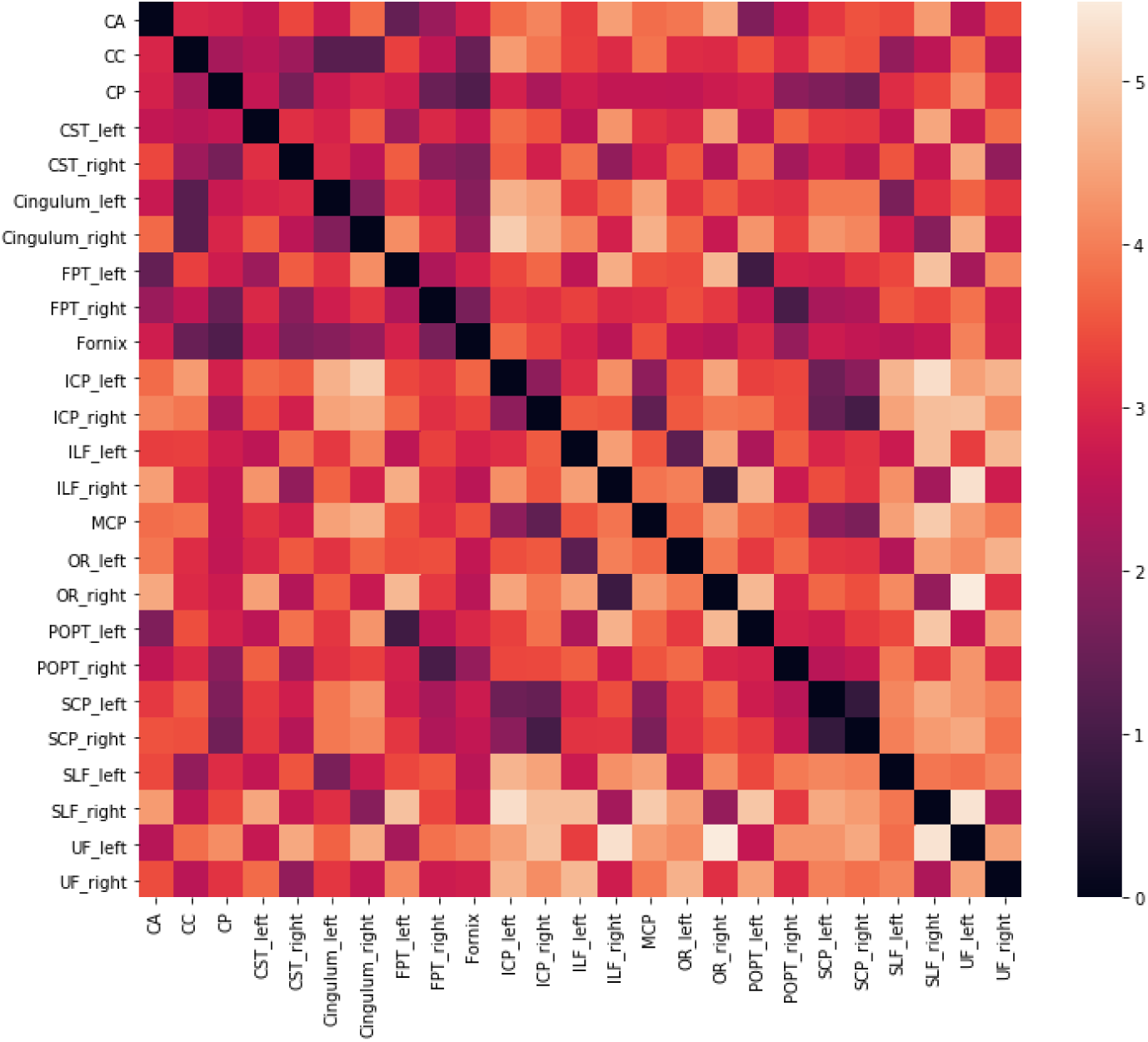
Bundle-wise distance matrix. Each bundle is represented by its bundle embedding which is the average of all the streamlines in this bundle. The bundle-wise distance matrix is constructed by computed the pairwise distances among all the bundle types.

### 3.5 Hierarchical Streamline Clustering and Streamline Filtering

Fig 9 shows a hierarchical dissection of the streamlines in the corpus callosum using unsupervised clustering of the streamline embedding. It is similar to a binary tree and each non-leaf node is clustered into two groups. It can be seen that the dissection was along the anterior and posterior direction and the iterative process segmented the CC bundle into two, four and eight clusters on level 1, 2 and 3 respectively. In this case, we used two clusters for each non-leaf node although any number of clusters can be set, and different numbers of clusters can be used for non-leaf nodes on the same level. For example, on level 1, the first sub-bundle can be further segmented into three sub-parts, while the second sub-bundle can be segmented into four sub-parts.

**Fig 9.**
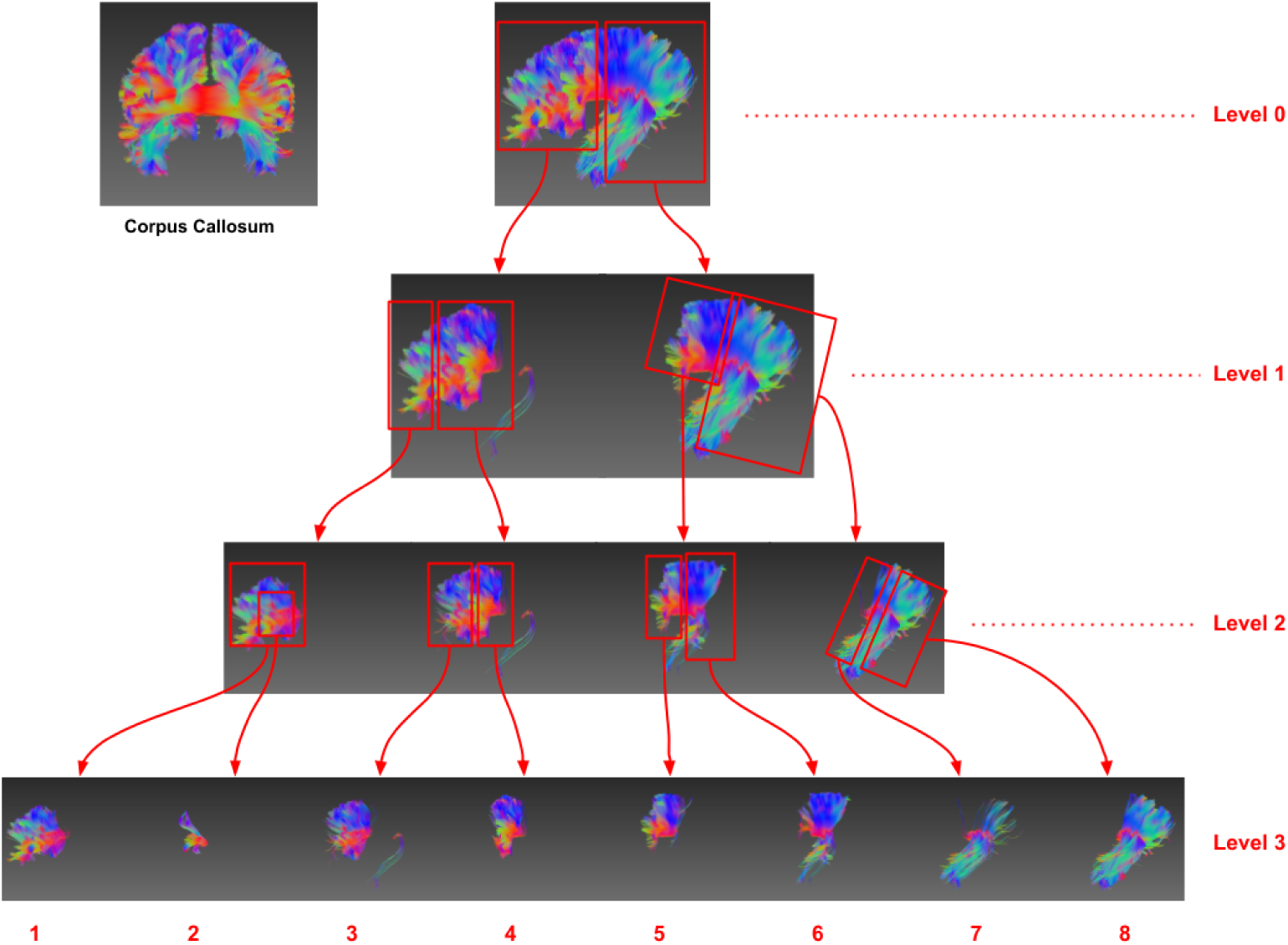
Hierarchical dissection of streamlines in the corpus callosum using streamline embedding clustering. On each level, the sub-bundles are further clustered into two groups iteratively. On level 3, the CC bundle can be dissected into eight sub-bundles along the anterior-posterior direction. Among the sub-bundles on level 3, the second leaf sub-bundles are identified as false streamlines.

Unsupervised clustering of streamline embedding can be used to identify and remove false streamlines. In Fig 9, the first sub-bundle on level 2 was further segmented into two parts (i.e. the sub-bundle 1 and 2 on level 3), with the streamlines in the second sub-bundle on level 3 identified as false connections (in this case, the streamlines seeded from cortical region in the left hemisphere, had a sharp turn in the past in the right hemisphere and ended in the seeding region). This finding highlights that the proposed technique can filter false streamlines generated in the tractography process as embedded streamline are spatially separable in the latent space.

## 4. Discussion

We present a recurrent neural network to encode streamlines with variable lengths into fixed-size latent vector representations and quantify similarities by measuring the Euclidean distance between the streamlines. Streamline bundles were represented as the average of the streamline latent vectors. With both embedded streamlines and bundles in the same vector length, the method enables quantitative analysis of streamlines at any granularity of streamline bundles. We validated the method using the ISMRM 2015 tractography challenge dataset, and showed that embedding streamlines to a learnable latent space enables efficient similarity measurement and streamline clustering.

The method is flexible such that any streamline length can be embedded into a latent vector of variable size. Existing methods generally resample streamlines to ensure they have the same length as required for streamline-wise distance measurements (Garyfallidis et al., 2012) and as inputs into neural networks for classification (Colon-Perez et al., 2016; Liu et al., 2019). CNN-based auto-encoders suffer from the same length limitation due to the fixed input size of CNN (Lam et al., 2018). However, the resampling process results in the loss of detailed information of the streamline and treats long and short streamlines differently. Our RNN-based method naturally supports variable lengths of sequence input without the need to truncate or resample the original streamlines. Unlike manually designed features (Yendiki et al., 2011; Berto et al., 2021), the latent space dimension in the proposed framework can be customized by adjusting the RNN hidden dimension. RNN networks with larger hidden dimension, i.e. higher dimension of the latent space, will have better capability of representing longer and more complex streamline structures, model larger numbers of streamlines, while smaller latent spaces require less computation. Hence, the size of the latent vectors is flexible and can be tailored in a case-by-case manner. In our work, a 128 dimension latent space was used, which was effective for modelling the streamlines in the ISMRM 2015 tractography dataset.

Sequential information is important when encoding streamlines into latent representations. A streamline is essentially a sequence of 3D spatial points that represent its location and shape in the brain. Treating a streamline as a collection of points may lose its sequential information during embedding. Both distance-based and even recent deep learning methods that are based on CNN networks fail to capture the sequential components. CNN network architectures are designed to be translationally invariant, and do not view the streamlines as an ordered sequence but instead process it, for the case of auto-encoding streamlines, as a 1D image with three channels. CNNs are sensitive to streamline segment patterns but ignore positions in the streamlines. Unlike CNNs, RNNs not only treat sequences naturally but provide a better representation capability for long sequences, i.e. streamlines with long length. Adding position encoding techniques to CNNs may compensate for the weakness of dealing with sequential data, similar to recently introduced techniques such as the Transformer architecture that could potentially be used for encoding sequential data types with attention mechanisms (Wolf et al., 2020). However, such approaches are not the focus of the current paper and will be explored in future work.

The latent space learnt with RNN auto-encoding provides an efficient way to form streamline bundle representations using data-driven clustering algorithms. Any granularities of the streamline bundle can be represented using the average of the embedded streamlines in the bundle in a computationally efficient manner. Distances can be measured between streamlines and also between a streamline and a bundle, since they are in the same closed latent space. The framework represents each streamline and bundle as tabular data, opening the possibility of analysis with generic data mining algorithms (e.g. anomaly detection) capable of dealing with high dimensional data. Bundle embedding can also be used to identify group-level landmarks (Gori et al., 2016) and create a multi-subject atlas (Guevara et al., 2012) in a data-driven and unsupervised manner. Finally, using the trained decoder the embedded bundle can be projected back to the 3D space for visual inspection purposes.

Of crucial importance is whether the streamline groups or bundles generated from unsupervised clustering algorithms in a latent space are anatomically plausible. We validated the streamline groups obtained using k-means clustering on the left brain hemisphere and manually examined each group, identified their similarity to the ground truth bundle types and found that the major bundles were extracted with minimal bias. Our motivation was not to re-discover well defined bundles, but rather to explore smaller structures, streamline filtering and hierarchical dissection. Qualitatively, the method requires a human expert to visually inspect the identified sub-bundle structures and determine their anatomical validity. The framework provides a data-driven streamline parcellation tool to study sub-bundle structure as an alternative to traditional cortical atlas based methods (Sporns et al., 2005; Bullmore & Sporns, 2009). Quantitatively, the method has the potential to be used with graph theory analysis over populations (Yeh et al., 2016; Bassett & Bullmore, 2017) to study connectivity patterns at various granularities of streamline parcellation.

The performance of the method can be improved by encoding streamlines from both directions and further validated using non-synthetic datasets. In the current work, the encoding was initiated from one direction whereas bidirectional LSTM (biLSTM) views of streamline in the auto-encoding process may improve the robustness of the embedding model. Alternatively, the effectiveness of using the Transformer architecture could be explored to learn a latent space to represent streamlines. The method was validated on the ISMRM 2015 tractography challenge dataset. Further validation work using other publicly available annotated datasets will be explored in future work.

One of the significant potential applications of the technique is automatic classification of streamlines for a human tractogram based on the distance of streamlines to the predefined bundle types in the training dataset. The training dataset in the current work had 25 predefined bundles types each represented by an embedding vector. Tractograms generated on other datasets, e.g. Human Connectome Project (HCP) dataset (Van Essen et al., 2013), could be embedded into the latent space using the trained auto-encoder with each streamline classified by its nearest distance measure to the 25 bundle types. The distance metric could then be used to determine a confidence interval with smaller distances indicating high confidence and vice versa. Such an approach could potentially offer an efficient classification solution to connectivity analyses in large scale population datasets.

## 5. Conclusion

In this work, we proposed a novel recurrent neural network auto-encoder to embed any length of white matter streamlines and at any level of bundle granularity, to a fixed-size latent representation for parcellation of whole-brain tractograms. The embedding process preserves the position, shape and sequential information of the streamline and bundle. The method was validated on a synthetic dataset and distinguished symmetric bundle structures, produced bundle groups with unsupervised clustering algorithms, and achieved good classification accuracy based on distance measurements on an annotated dataset. The method can be applied to query streamlines according to similarity, hierarchically dissect streamline bundles for studying sub-structures, and identify and remove biased streamlines. The method enables the possibility of quantitative distance analysis using data mining and unsupervised learning techniques.

## Information Sharing Statements

The source code of the proposed work is freely available for non-commercial use from https://github.com/BioMedAnalysis/latenttrack.

